# No evidence for general intelligence in a fish

**DOI:** 10.1101/2021.01.08.425841

**Authors:** Mélisande Aellen, Judith M. Burkart, Redouan Bshary

## Abstract

Differences in human general intelligence or reasoning ability can be quantified with the psychometric factor *g*, because individual performance across cognitive tasks is positively correlated. *g* also emerges in mammals and birds, is correlated with brain size and may similarly reflect general reasoning ability and behavioural flexibility in these species. To exclude the alternative that these positive cross-correlations may merely reflect the general biological quality of an organism or an inevitable by-product of having brains it is paramount to provide solid evidence for the absence of *g* in at least some species. Here, we show that wild-caught cleaner fish *Labroides dimidiatus*, a fish species otherwise known for its highly sophisticated social behaviour, completely lacks *g* when tested on ecologically non-relevant tasks. Moreover, performance in these experiments was not or negatively correlated with an ecologically relevant task, and in none of the tasks did fish caught from a high population density site outperform fish from a low-density site. *g* is thus unlikely a default result of how brains are designed, and not an automatic consequence of variation in social complexity. Rather, the results may reflect that *g* requires a minimal brain size, and thus explain the conundrum why the average mammal or bird has a roughly 10 times larger brain relative to body size than ectotherms. Ectotherm brains and cognition may therefore be organized in fundamentally different ways compared to endotherms.

## Introduction

Fish do many apparently smart things (1,2). Yet, being ectotherm vertebrates, they have on average ten times smaller brains corrected for body size compared to endotherm vertebrates (3,4). Size differences are even more pronounced if only the pallial part of the forebrain is considered, i.e. the part that is responsible for more complex cognitive functions (5,6). So what can mammals and birds do with big, physiologically expensive brains (7,8) that reptiles, amphibians and fishes cannot? A possible hypothesis is that only endotherms have general intelligence. In humans, general intelligence in its broad definition involves various domains of cognition, such as reasoning, planning, problem-solving as well as learning from experience, that are centrally integrated to allow for more flexible behaviours, to understand abstract concepts, and to have conceptual thoughts (9). A good indicator of the presence of general intelligence in humans is the psychometric factor *g. g* results from the positive manifold, i.e. the well-established finding that in humans, individual performance across tasks testing different domains is correlated (10–15). Factor analyses of performance across such tasks will thus result in a first factor on which all tasks load positively, and this general factor is referred to as *g. g* has been demonstrated for a variety of mammals in controlled laboratory experiments, including average brained species such as mice (16). Evidence for *g* is less clear in birds (17) (but see (18)) but those studies have been conducted on small numbers and/or in the field, potentially creating biased data as mostly motivated and/or bold individuals will participate under these conditions (17).

Using *g* as indicator of general intelligence species other than humans has been criticised with the argument that the positive manifold might be a pure side-effect. For instance, the positive manifold may simply reflect variation in low-level biological properties, due to ontogenetic disturbances, or genetic load (i.e. the accumulation of deleterious, pleiotropic mutations, e.g. (19,20)). Individuals with less disturbances, or less genetic load, may more fully express their growth potential, which may also lead to better myelination of the nervous system, which ultimately will operate smoother and faster across domains (21). Moreover, a positive manifold can be an artefact of how brains are generally organized. Thomson (22) pointed out already in 1916 that a positive manifold can arise in the absence of general intelligence due to between-task neural overlap (see also (19,23–28)).

Thus, our aim was to simultaneously test two closely linked hypotheses by using cleaner fish *Labroides dimidiatus* as study species. First, is general intelligence restricted to endotherms? Second, does a *g* – factor mandatorily arise whenever a large enough, unbiased sample of animals is tested under appropriate conditions? *L. dimidiatus* is a particularly suitable study species as there is plenty of evidence suggesting that its cognitive performance is rather outstanding for an ectotherm vertebrate. It engages in interactions with client fish that visit to have ectoparasites removed while cleaners prefer to eat client mucus (29). Most likely due to this conflict of interest, cleaners show high strategic sophistication in ecologically relevant tasks, even outperforming primates (30). Furthermore, cleaners are able of generalised rule learning (31), and apparently even pass the mirror test (32). Moreover, as variation in intraspecific social complexity affects performance in *g* tasks in birds (18) and as it is well established that fish densities affect cleaner fish expression of strategic sophistication in ecologically relevant social tasks (33,34), we compared 80 individuals from two sites. One site harbours high cleaner densities and the other one low densities (3.3 versus 0.4 cleaners / 100m^2^, see SI, S2 Fig), which correlate strongly with large client densities (33). If cleaners from the high-density site perform better also in *g* tasks, this would increase the likelihood that we would find a positive manifold. Taken together, according to current knowledge we maximised the chance to find *g* in an ectotherm vertebrate, which would support the notion that *g* is an artefact and refute the hypothesis that *g* requires large brains.

## Results

We conducted four laboratory experiments (Fig 4) that have also been used on mammals before (16). The four experiments tested for different cognitive domains: flexibility (reversal learning; abbreviated ‘RL’), self-control (detour task, abbreviated ‘DT’), numerical competence (dot number task, abbreviated ‘NC’) and working memory (object permanence). As tests for *g* have to warrant performance above chance without ceiling effects, we had to omit the object permanence task (presented in SI, S5 Fig) but could keep the other three tasks. Of the 80 subjects, we had to remove eleven individuals because they did not participate in at least one task. With the remaining 69 individuals, we used a principal component analysis to test for the presence of a positive manifold (Fig 1). Dimension 1 explained 38.5 % of the variance in performance, with an Eigenvalue of 1.16. Two tasks loaded positively (RL and DT), whereas the other one loaded negatively (NC). Dimensions 2 and 3 explained rather similar amounts of variance (32.8 and 28.7 %; Eigenvalues 0.98 and 0.86). The results thus revealed no evidence of a psychometric *g* in *L. dimidiatus*. Indeed, correlations of individual performance across tasks support the view that the performances in the three tasks are independent of each other (Spearman-Rank correlations, all n = 69, RL-DT: *r* = 0.026; RL-NC: *r* = −0.137; DT-NC: *r* = −0.051; all NS, Fig 2a-c). Also, separate analyses for cleaners from high- and low-density sites did not reveal any evidence for *g* (SI, S3 Fig).

**Fig 1.**
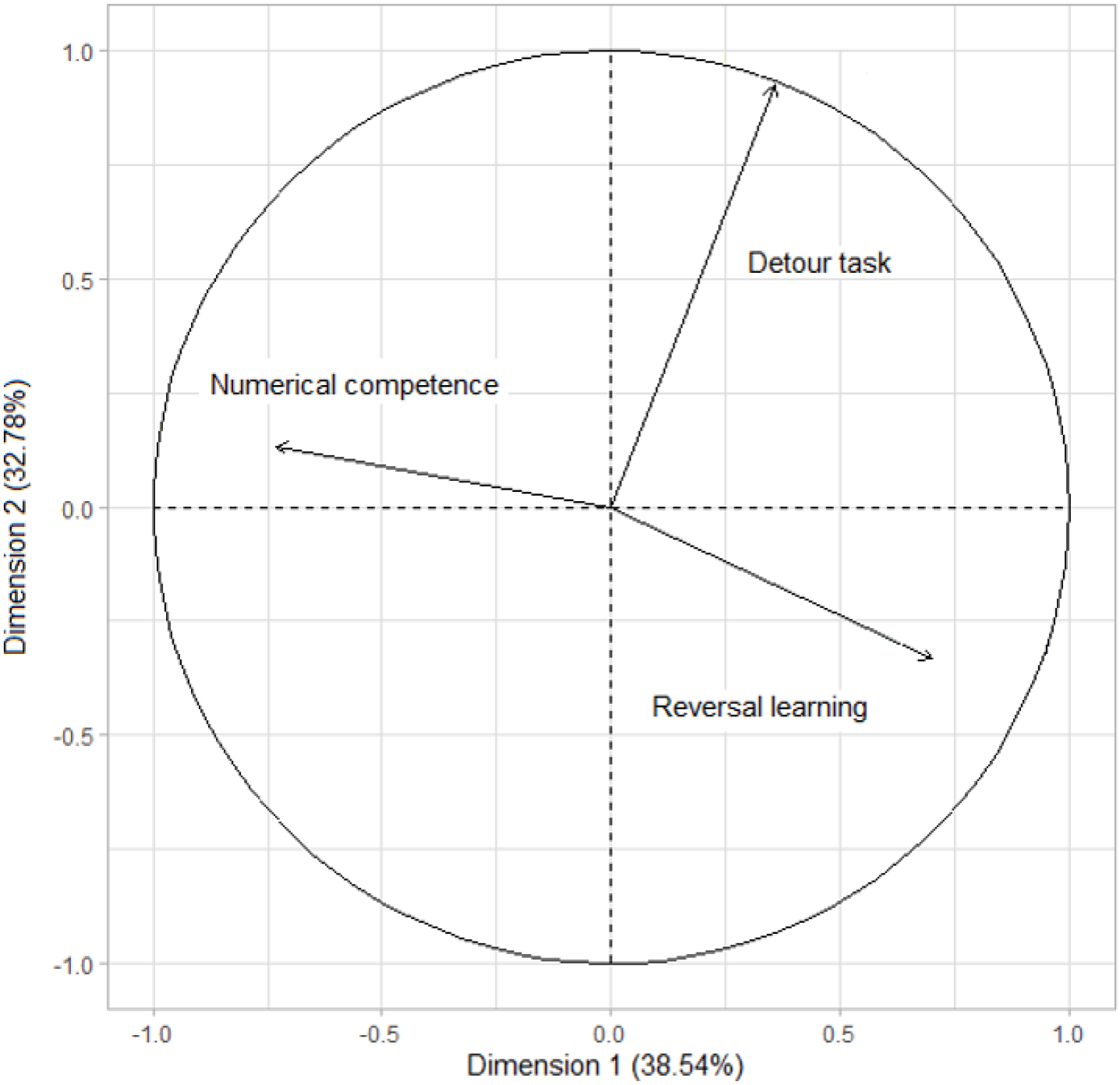
No evidence for *g* in a fish. Principal Component Analysis (PCA) of the three cognitive tasks. Dimension 1 (explaining 38.54 % of the variance in performance) and dimension 2 (explaining 32.78 % of the variance) are represented. The results for each task are represented as vectors.

**Fig 2.**
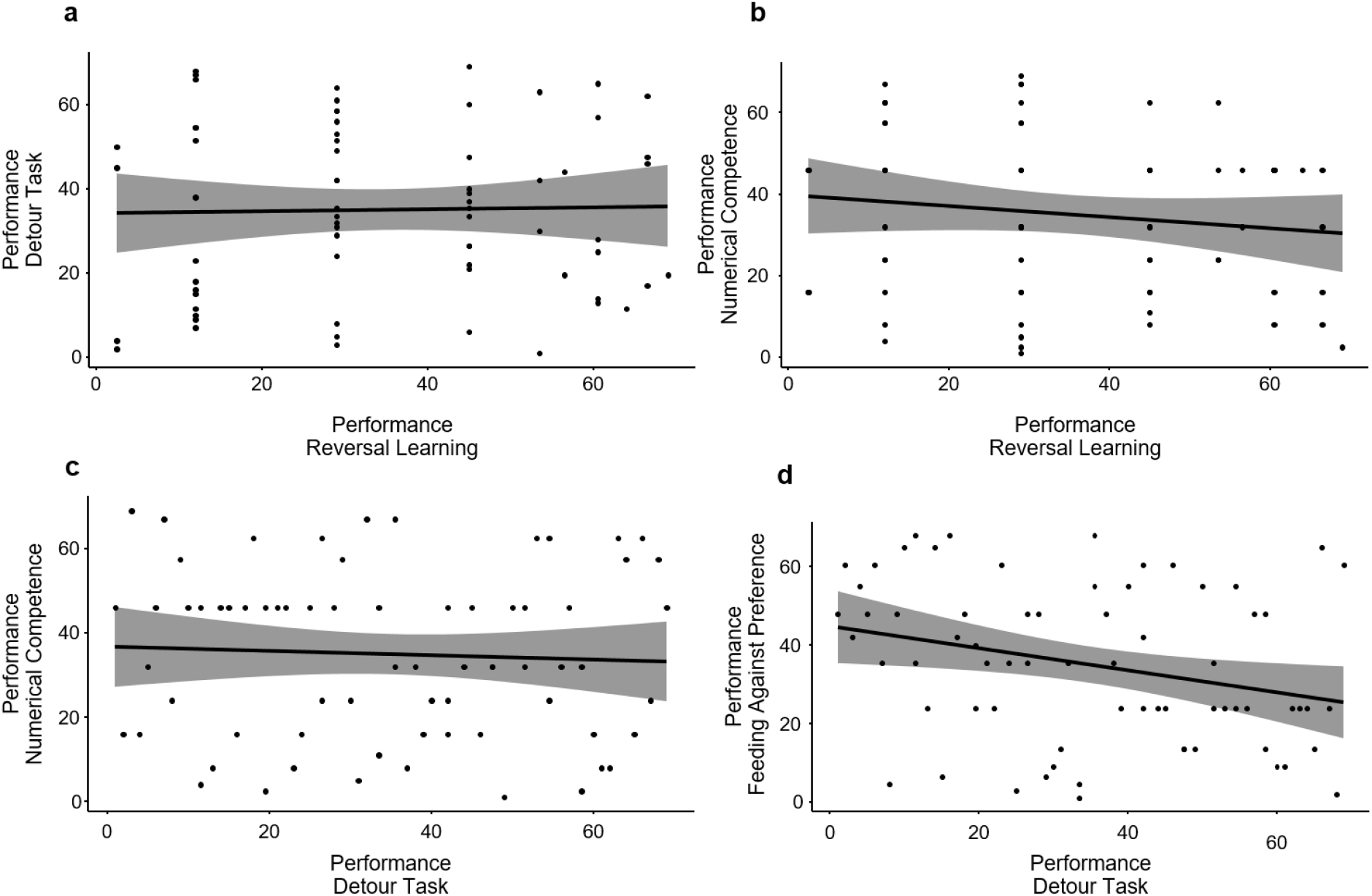
Lack of positive correlations across cognitive tasks. Pairwise correlations between the ranks of individual performances in four different cognitive tasks (rank 1 being highest performance, rank 69 being lowest performance). Individual performances did not significantly correlate between any two of the three g tasks (a) RL – DT, Spearman-Rho r = 0.026, p = 0.8; b) RL – NC, Spearman-Rho r = −0.137, p = 0.3; c) DT – NC, Spearman-Rho r = −0.051, p = 0.7). The only significant correlation was negative rather than positive, and found between DT and FAP (d) Spearman-Rho r = −0.283, p = 0.02 (p = 0.03 with Bonferroni correction)), a *g* task and an ecologically relevant task that are both testing for inhibition.

In addition to the *g* tasks, we conducted one ecologically relevant experiment, i.e. the individuals’ ability to feed against preference (abbreviated FAP) in order to prolong interactions with clients and hence obtain more food. Like the detour task, the experiment measures self-control, allowing us to ask how performance in an ecological task relates to performance in an abstract task within a single domain. We found that DT was negatively correlated with FAP (Spearman-Rho *r* = −0.283, p = 0.018, Fig 2d). While not expected, the result suggests that positive correlations in performance may even be absent within domains. Comparisons of cleaners from high- and low-density sites revealed no consistent differences in the performance in each task separately (Fig 3). Differences were far from being significant for all tasks (RL (panel a), DT (panel b), NC (panel c), and FAP (panel d) (Wilcoxon-tests, all p > 0.13). There was also no difference in cleaner body length between sites (Wilcoxon-test, p = 0.24; SI, S4 Fig), suggesting no systematic differences in age that could potentially have confounded the results.

**Fig 3.**
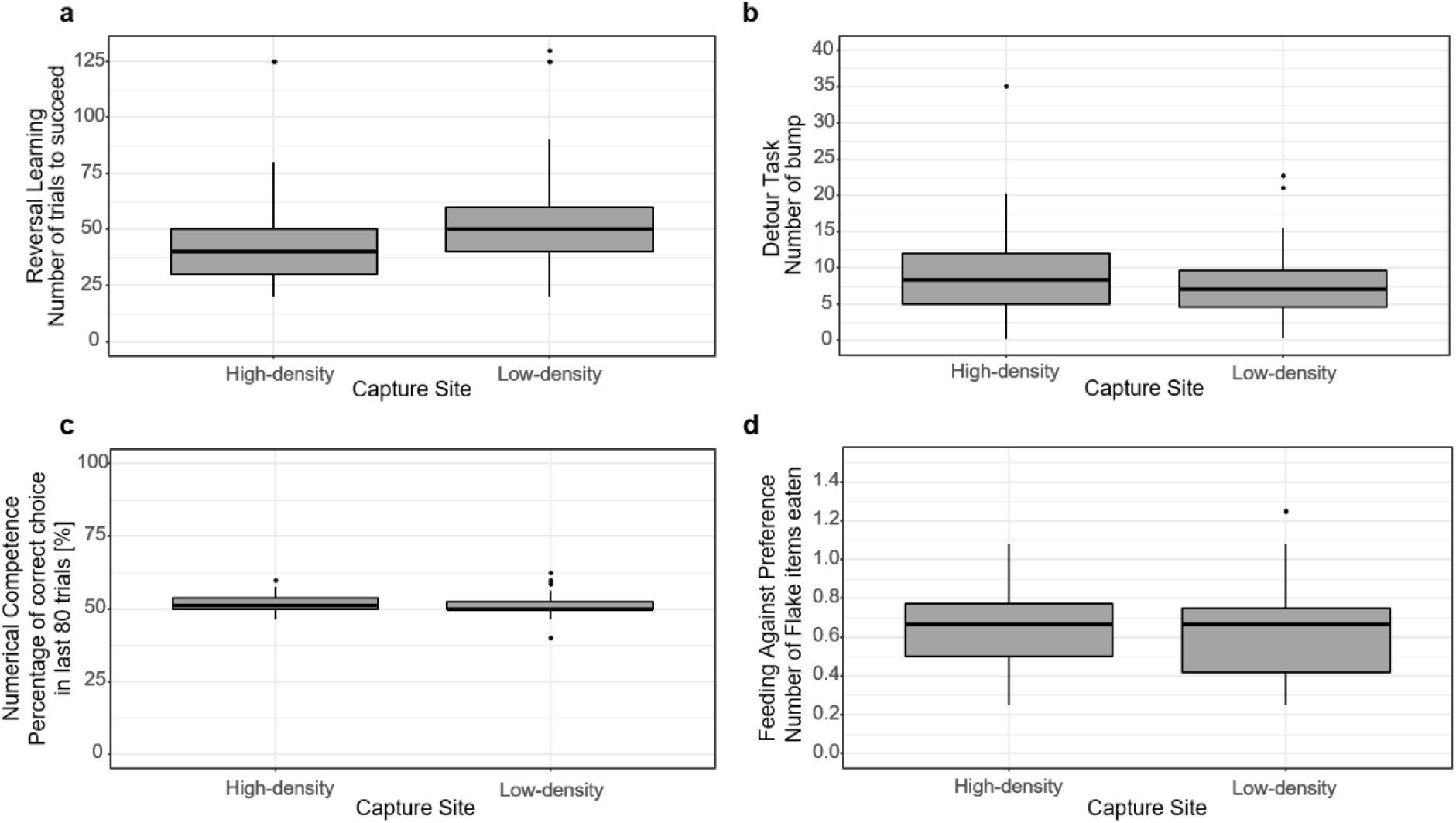
Fish densities do not affect performance. Box plots showing median, interquartiles and 95 % range of individual performances split by site of capture (high-density and low-density site). a: number of trials to complete reversal learning task; b: mean number of head bumps into Plexiglas separation in the detour task; c: % correct choices in the numerical competence task; d: mean number of flakes eaten before a prawn item (i.e. highly preferred food) in the feeding against preference task.

## Discussion

The lack of correlated performance, and thus *g* in this fish species has far-reaching implications. First, given that we tested a large random sample of cleaner fish under controlled laboratory conditions, we provide the strongest evidence as yet that a positive manifold does not emerge as a default in brains. Instead, our results support the notion that a major consequence of the differences in brain size between mammals and fish is that mammals have some system-level cognitive abilities that can be used across tasks tapping into different cognitive domains, while the fish brain is organised in a more modular way. A more modular organisation does apparently not prevent the emergence of complex cognitive processes like transitive inference, generalised rule learning and even mirror self-recognition (31,32,35), i.e. processes that go beyond conditioning. Thus, a presence/absence of these cognitive processes cannot explain the ten-fold difference in relative brain size and the even larger difference with respect to the pallial brain part (5,6) between endotherm and ectotherm vertebrate clades. Also, the cleaners’ performance in at least two *g* tasks (reversal learning and detour) was very good, and there is evidence for high numerical competence as well as object permanence in other fishes (2,35), suggesting that single cognitive tasks used to test for general intelligence are not per se too difficult for ectotherms. In contrast, the transition from a modular organisation to more general-purpose intelligence may have required a massive increase in brain size.

A second major insight from our results is that environmental complexity, as indicated by cleaner and other fish densities, largely fails to affect performance of cleaner fish in *g* tasks, in contrast to results from Australian magpies (18). Therefore, it appears that previously documented differences in strategic sophistication in ecologically relevant tasks between cleaner fish individuals from high-density and low-density sites (33,34) are due to specific experience effects rather than cognitive abilities per se.

Currently, most discussions on brain evolution focus on the relative importance of social versus environmental complexity. While this discussion remains key to explain variation in size and structure within clades, the fundamental divide between endotherm and ectotherm vertebrates brain sizes may be largely driven by brains being more general-purpose machines in the former versus specialised learning machines in the latter.

## Materials and Methods

### Cleaner fish: *Labroides dimidiatus*

The cleaner wrasse, *Labroides dimidiatus*, is a protogynous fish and lives in a small territory called cleaning station (36). It lives in harems and males can have up to five females in their territory comprising several cleaning stations (one per female) (37). The species is widespread in the Indopacific ocean and can also be found in the Red Sea (38). It feeds on the surface of other reef fish called clients by removing ectoparasites from them. Cleaner fish have around 2000 interactions per day (39). As cleaner fish prefer to eat mucus over ectoparasites (29) this creates a conflict of interest between cleaner and client over what the cleaner should eat. As a consequence, cleaners need to eat against preference in order to cooperate, and hence to avoid that clients respond with evasive actions like chasing the cleaner or switching to a different partner for their next interaction (40). Therefore, the experiment in which cleaners needed to feed against preference reflects high ecological relevance (41).

### Capture, individuals housing, and acclimation

The study was conducted at the Lizard Island Research Station, Great Barrier Reef, Australia in February - May 2018 and 2019. By finding pairs of cleaners and avoiding the larger individual, 80 female cleaner fish were caught with a barrier net (2 m long, 1.5 m high, mesh size 0.5 cm; 40 fish each year) and hand-nets on nearby reefs. 40 individuals were from a high-density client fish area (Birds Islet crest; 20 each year; SI, S1 Fig), and another forty were from a low-density client fish area (Birds Islet lagoon; 20 each year; SI, S1 Fig). 30 m long transects covering a width of 5m (for detailed methods see (42)) revealed that mean cleaner fish density, which is highly correlated with the density of large clients (33), differed by a factor 8 (3.3 versus 0.4 individuals / 100m^2^; Wilcoxon-test, m = 6, n = 5; p = 0.012; SI, S2 Fig). Fish were housed individually in glass aquaria (62 x 27 x 38 cm). Each year, the forty individuals were caught at the beginning of the field trip and split into two experimental cohorts of 20 individuals each, which were tested simultaneously. Cleaner fish were acclimatized for at least twelve days before being subjected to five different experiments (experimental cohorts 1 and 3) or for at least 44 days (experimental cohorts 2 and 4). Cleaner fish were acclimated to feed on Plexiglas plates, mimicking client fish in the captive environment. We provided mashed prawn as food that we smeared on the Plexiglas plate. When fish were well accustomed to their feeding plate (which took two to three days), they were trained to eat small pieces of mashed prawn placed on dots drawn on a new feeding plate. Once fish were eating invariably well on the feeding plate with and without dots, they were habituated to the different plates and barriers that we used during all the different experiments (Fig 4). Most notably, fish were exposed to a barrier that divided the aquarium into the holding compartment and the experimental compartment. A door (dimension 7 x 18 cm) in the barrier could be opened so that the cleaner fish could swim through. Cleaners thus had to be habituated to swimming through the door.

**Fig 4.**
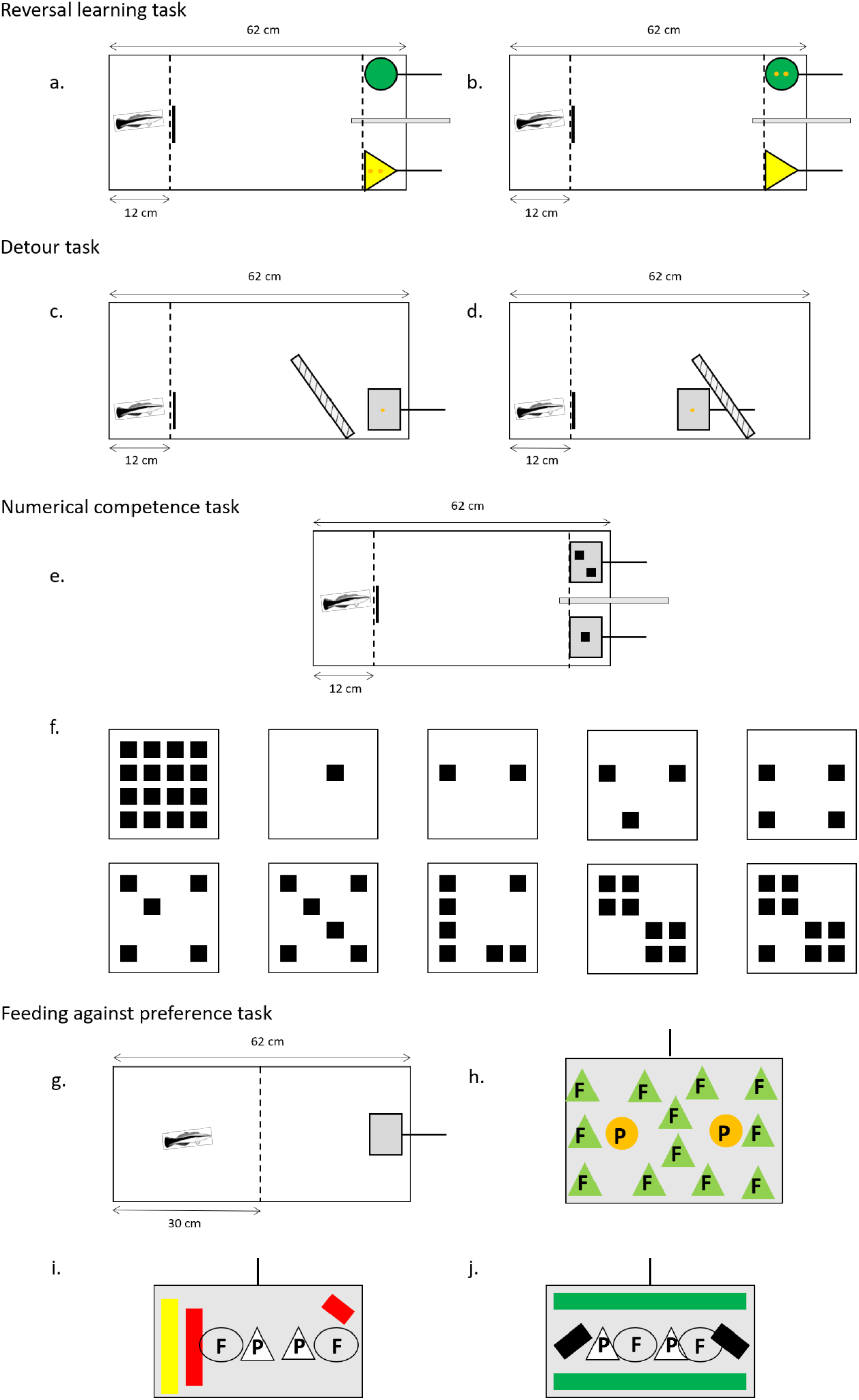
Spatially explicit experimental setups of the three *g* tasks (a – f), and experimental setup of the ecologically relevant task, the ability to feed against preference in order to increase food intake (g – j). Dashed lines: the transparent barrier that separated the holding compartment from the experimental compartment (with the short black line indicating the door through which fish could cross). Thicker grey and hatched structures represent opaque and transparent barriers. Orange dots on the plates show the location of food rewards (mashed prawn items). Panel a. shows the initial associative learning task, upon its completion we started the reversal learning task by changing the role of the two plates (panel b.). Panels c. and d. show the positions of the reward plate in the detour task, where only trials with the reward plate behind the barrier were analysed. Panel e. shows the experimental setup for the numerical competence task. In panel f., the plate to the upper left indicates the 16 possible positions for the back squares, while the other plates give examples for plates displaying 1-9 squares. g. general setup. The dashed line shows the see-through barrier that kept fish in the holding compartment while the experimenter placed a plate with food items in the experimental compartment. h. the plate used to train fish that eating flake items is allowed while eating a prawn item leads to the immediate removal of the plate. F: flake item. P: prawn item. i. and j. the two plates used alternatingly in the actual experiment, offering 2 flake and 2 prawn items.

### General experimental procedures

During all experiments, subjects were guided into the holding compartment before each trial. This allowed the experimenter to subsequently to set up the trial (i.e. plates with or without food, partitions, barriers) without the fish intervening. Once a trial was properly set up, the door was opened so that the cleaner fish could swim through and make its choices. During simultaneous choice tasks (reversal learning an numerical competence tasks), an opaque separation (10 cm wide) between the two plates helped to define a subject’s choice: when the subject’s head passed the imagined line perpendicular to the start of the opaque separation (Fig 4a,b,e), we scored that the subject had chosen the plate inside the compartment. If the choice was wrong, the experimenter removed the correct plate immediately. If the choice was correct, we allowed subjects to still inspect the wrong plate after having finished eating the reward before removing both plates simultaneously.

In all tasks, the experimenter tested each fish once before moving on to the next fish. Once each fish had been exposed to one trial the next round of trials began, leading to an intertrial interval of about 30 min. Twenty trials could be completed within a day, starting at 6:00 and finishing at 17:00.

Fish conducted the experiments in their home aquarium. All experiments were video recorded with a GoPro mounted on the forehead of the experimenter. All fish from 2018 and 18 fish from 2019 were released at their respective sites of capture after experiments had been completed. Another 22 fish were used in another project that included brain analyses. The project was approved by the Animal Ethics Committee of the Queensland government (DAFF; AEC Application Reference Number CA 2018/01/1155).

Before and in between the experiments, the fish were fed ad libitum every day, introducing the feeding plate in the morning and removing it at the end of the day. During experimental days, fish had to obtain food from making the correct choices as long as trials were conducted. They were fed ad libitum at the end of the day after the trials had been completed. One day off was kept between each experiment.

#### Experimental setups

### Specific experimental protocols

#### 1. Reversal learning task: learning – flexibility (Fig 4a,b)

In reversal learning tasks, subjects first learn to associate one of two stimuli with a reward. After this initial association learning, the contingencies are reversed and the other stimulus is rewarded. This task thus measures how flexibly individuals can adjust to the new reward contingencies. For this task, we offered two plates simultaneously: a yellow triangle shape Plexiglas plate (8 cm wide and 7.5 cm height) was placed on the left side of the aquarium with two pieces of prawn located on the back. On the other side, a green round shape Plexiglas plate (8 cm wide and 7.5 cm height) without food was positioned. The positions of the two plates remained constant during a first training phase, testing for initial learning and during the reversal learning phase. For each trial, fish were first placed behind a transparent grid barrier with a transparent door on one side of the aquarium, 12 cm from the aquarium wall to form the waiting area. The plates were then introduced on the other side (with the handle leaning against the aquarium wall providing stability), and only then the door was lifted and the fish could make its choice by approaching one of the 2 plates.

To facilitate learning the initial association, we put two pieces of prawn on the back of the rewarding plate and left both plates in the aquarium. Initially, individuals different with respect to how fast they would approach both plates. We hence varied the duration of first trials such that all individuals would inspect both plates and eat the food, yielding trial durations between five and 30 min. within two days, trial durations were down to 30 – 60 seconds. Such time intervals allowed the fish to eat the two items and to confirm that the other plate did not offer any food. This training phase consisted of two exposures per day over ten consecutive days. We then tested the fish for a significant preference for the rewarding plate, offering only one item per trial, conducting 2 sessions of ten trials each per day. The criteria for success were either 10/10 or 9/10 correct choices in a session, twice 8/10 in two consecutive sessions, or at least 7/10 in three consecutive sessions. Once an individual had reached criteria for the initial preference, we reversed the role of the two plates. Thus, two pieces of mashed prawn were now placed behind the green round shape Plexiglas plate on the right side of the aquarium. Each fish performed 20 trials per day and was tested the same way according to the same learning criteria as described for the initial learning. We initially ran trials for up to five days. If a subject had not succeeded yet within these 100 trials, we conducted an extra five trials in which we prevented subjects to swim to the yellow plate by inserting a see-through barrier in front of it. These extra trials either ensured that the individuals were exposed to feeding off the green round plate, or they showed that some individuals simply refused to approach that plate. We considered that these latter individuals were rather afraid of the plate and hence their performance could not be interpreted as a failure to learn. We therefore removed them from the data set. In contrast, we exposed the subjects that had eaten off the green round plate to another 20 trials on the 6^th^ day to see if after these extra five trials they could reach the criterion or not. The number of trials needed to reach learning criterion in this reversal learning task was used as measure of behavioural flexibility for the statistical analysis.

#### 2. Detour task: inhibitory control (Fig 4c,d)

We tested if cleaners were able to swim around an obstacle to get a food reward. This task permitted to measure inhibitory control as well as spatial problem-solving. On the day prior to the first test we familiarised subjects with the anthracite Plexiglas plate (10 cm x 5 cm) that offered a visible piece of mashed prawn on its front side. We also acclimated the fish to the obstacle by inserting it in the aquarium for 45 minutes twice during the day (once on the right-hand side and once on the left-hand side). Given that cleaners had never been tested in a detour task, we did not know what level of performance we could expect. We therefore decided a priori to start with a simple task and to increase difficulty on consecutive days as long as subjects readily managed to access the food reward. On the first day, we placed a transparent obstacle in form of a plate, made visible by drawing a grid of black lines (1 – 1.5 cm apart) onto it, in front of a food plate. The obstacle was placed perpendicular to the aquarium side wall. It was 19 cm wide, leaving subjects 5 cm to swim round it to get to the food plate 15 com behind the obstacle during test trials. In 50% of trials, the reward plate was placed in front of the obstacle. During the total of 10 morning trials, the plate, obstacle and door were always on the right side from the cleaners’ perspective. In the 10 afternoon trials, all equipment was moved to the left side from the cleaners’ perspective. As cleaners accessed the food plate without any problems, we counterbalanced the position of equipment within sessions on the second day, conducting no more than two consecutive trials on any side. As subjects continued to perform well, we placed the obstacle at a 45 degrees angle (Fig 4c,d), further increasing the inhibition requirement because fish had to swim away from the food in order to access it. We only used data from day 3 for our analyses. Subjects had a maximum of 60 secs to reach the reward plate. While we measured time to complete a trial, that measure may not well reflect cognitive performance as it is influenced by swimming speed, which may vary according to body size and/or motivation. Instead, we recorded the number of head bumps against the obstacle in each trial as a measure of a cleaner’s ability of self-control. Only trials in which the plate was behind the obstacle were analysed; trials in which the plate was in front of the obstacle only served to prevent the development of route routines. A total of 20 trials (10 experimental and 10 controls) were conducted, with an inter-trial interval of about 30 min.

#### 3. Numerical competence task: quantitative reasoning (Fig 4e,f)

This task tested quantitative reasoning in fish in a general form. In each trial, subjects were presented two white plates (7.4 cm x 7.4 cm) with differing numbers of black squares on them (Fig 4e). Each square was 11 mm^2^ in size, and the number of squares on a plate varied between 1 to 9. On the day prior to the experiment, we acclimated the fish to the plate first by smearing mashed prawn on a version displaying 10 squares. We later conducted two presentations where we placed 2 prawn items on the back of the plate. During experiments, we used in total 20 different combinations of square numbers: 5:1, 6:3, 6:2, 6:4, 4:3, 3:2, 2:1, 4:1, 5:2, 3:1, 5:3, 7:3, 7:4, 7:2, 9:3, 8:3, 8:4, 8:5, 9:4, and 9:5. Each combination was presented once per day, over a total of 8 days (yielding 160 trials in total), with the order of presentation randomised between days. There were in total 16 potential positions for a square on a plate (Fig 4f). We randomised the positions of squares for each number, removing any configurational cues for cleaners to make choices. Across the 8 days, the position (left or right) of the two plates in each combination was counterbalanced. The plate with the greater number of squares invariably offered two food items on its back, while the other plate contained inaccessible food on its back. Thus, some plates (those with 2-6 squares) sometimes yielded food and sometimes they did not, depending on which plate they were paired with. As a consequence, only the learning of a general rule based on numeric competences would yield performance above chance levels (“always choose the plate that shows more squares”). Previous research on cleaner fish has shown that this species has numerical competence, being able to learn to prefer one plate over another one based on the number of black squares (rather than spread or total black surface area) (43).

#### 4. Feeding against preference task: inhibitory control (ecologically relevant task) (Fig 4g-j)

The first three tasks presented cognitive challenges in abstract experimental setups, testing for cognitive skills in the absence of ecologically relevant contexts. In contrast, the ability of feeding against preference is of high ecological relevance. In natural interactions with client fish, cleaners need to largely feed on less preferred ectoparasites (gnathiid isopodes) instead of preferred client mucus so that it pays clients to visit cleaners (29). Our task mimicked natural interactions, replacing clients, parasites and mucus with a plate, preferred prawn items and less preferred flake items. There were three flake and three prawn items on the plate. A trial continued as long as cleaners ate items, while the experimenter removed the plate as soon as a cleaner ate a preferred prawn item. The food-maximising strategy was hence to eat against preference. Therefore, the feeding against preference task is conceptually linked to the Detour task as it measures inhibitory control. We included the task to test whether there was any correlation between ecological and non-ecological tasks within a single cognitive domain.

In order to prepare the fish for the experiment, we first gave them fish flakes (a mixture of 20% fish flakes and 80% mashed prawn to make the food stick to plates) on feeding plates instead of pure mashed prawn to familiarise subjects with the new type of food. In the next step, we conducted six training trials using a 15 x 10 cm plate with 14 items displayed on it (12 flake and 2 prawn items (44); Fig 4h). During a single trial, the plate was removed for 30 s as soon as the cleaner ate the first prawn item, and then reintroduced to allow the cleaner a second feeding bout until it ate the second prawn item. With this design, all subjects ate flake items and could hence potentially learn that feeding on a prawn item leads to the removal of the plate, while eating a flake item has no negative consequences. The intertrial interval was of one hour.

The next day, we conducted the experiment. We exposed cleaners 12 times to a single plate (10 cm x 6 cm) with two flake and two prawn items (Fig 4i,j). For a different study on reputation management, we also conducted trials involving the simultaneous presentation of two plates, but those data were not relevant for the current analyses. We took the mean number of flake items eaten over the 12 trials as a measure of inhibitory control for the analysis.

### Data Analysis and Supplementary Results

In each experiment, we ranked each fish by its performance according to the criteria specified in each section above (SI, S1 Table). From the eighty fish caught for this study, we were able to run the analysis with 69 individuals. A total of four individuals died before completing all tasks, while seven failed to participate in at least one task. Of the 69 individuals, 36 were from the high-density site (Birds Islet crest), and 33 were from the low-density site (Birds Islet lagoon).

We first examined whether there was sufficient interindividual variation in the performance of the cleaners in each task using descriptive statistics (means with standard deviation, and maximum – minimum values; S1 Table). For the ND task, we also verified that cleaners as a group performed above chance using a non-parametrical Wilcoxon signed-rank test in IBM ^®^ SPPS ^®^ Statistics version 25.0.0.1 (Wilcoxon-test, p = 0.001). In all three *g* tasks, we found substantial variation in performance and neither floor nor ceiling effects so that the three data sets were included in a principal component analysis (PCA; using the packages FactoMineR and missMDA). The key question was whether individual performance in each task was loading positively on the first PC factor. In a second step, we calculated correlations in individual performance across all possible pairs of the three tasks, using Spearman-Rho correlations as presented in the main text.

In order to check the robustness of our results, we investigated whether non-cognitive variables such as the site of capture, the year, the experimental set, as well as body length could have had an effect on the variation found in the PCA using a linear model (package ‘lme4’) (45). Moreover, we analysed the individuals from the different capture sites separately. The PCA analysis with the individuals from the high-density site explained 41.1 % of the variance in performance in dimension 1, with an Eigenvalue of 1.23. Two tasks loaded positively (RL and DT), whereas NC loaded negatively. No significant correlations were found between the three different tasks (Spearman-Rank correlations, n = 36, RL-DT: *r* = 0.179; RL-NC: *r* = −0.086; DT-NC: *r* = −0.072; S3a Fig). In the PCA analysis with the individuals from the low-density site, dimension 1 explained 41.12 % of the variance in performance with an Eigenvalue of 1.23. Two tasks loaded positively (DT and NC), whereas RL loaded negatively. The correlations of individuals performance across tasks were invariable slightly negative and non-significant (Spearman-Rank correlations, n = 33, RL-DT: *r* = −0.095; RL-NC: *r* = −0.217; DT-NC: *r* = −0.010; S3b Fig). No difference was found in the body length between the high and the low-density captures sites (Wilcoxon-test, p = 0.12; S4 Fig).

To test the extent to which abstract and ecological tests yield similar performances, we correlated individual performances in the detour task and the feeding against preference task, using a Spearman-Rho correlation. Thus, we calculated in total four correlations with our data. As a consequence, we used Bonferroni correction to calculate a new α’ = 0.0125 (α/number of tasks = 0.05/4) to control for finding a spurious result. Finally, we used Wilcoxon-tests to evaluate whether performance in any one test differed systematically as a function of site of capture. Again, we used Bonferroni correction to control for multiple testing which resulted in an α’ = 0.0125 (α/number of tasks = 0.05/4. The Spearman-Rho correlation test, the PCA, the linear models as well as the Wilcoxon-Test were carried out in R Rstudio © (R Version 1.3.1093, © 2009-2019 RStudio, PBC).

### Data archiving

All data, meta-data as well as the R codes can be found on https://figshare.com/s/5ba3e10c1501ee1219ec.

## Acknowledgements

We are grateful to Lizard Island Research Station and their amazing staff. We thank in particular Carel van Schaik for stimulating discussions. We thank Yasmin Emery for her precious help collecting some data during the first two sets in the first year. We thank Radu Slobodeanu for statistical help and support. The study was supported by the Swiss National Science Foundation (grant number: 310030B_173334/1 to RB).

