## Supporting Information for "No evidence for general intelligence in a fish"

**S1 Table: Description of the tasks and their measurements.**

| Cognitive domain | Task | Performance measurement | Measurement description | Mean (SD) | Max – Min | N |
|---|---|---|---|---|---|---|
| Learning – flexibility | Reversal learning | Quantity | Number of trials needed to reach criterion | 51.02 (26.72) | 125 – 20 | 69 |
| Inhibitory control | Detour task | Quantity | Number of bumps done in 1 trial obstacle positioned at an angle | 8.49 (5.91) | 35.1 – 0.2 | 69 |
| Quantitative reasoning | Numerical competence | Percentage | Percentage of rewards retrieved in the 80 last trials | 51.54 * (3.67) | 62.5 – 40 | 69 |
| Inhibitory control | Feeding against preference | Quantity | Number of flake items eaten prior prawn in 1 trial 1 plate exposition | 0.65 (0.22) | 1.25 – 0.25 | 69 |

The mean, maximum and minimum values are from the 69 individuals that were included in the PCA* indicates the performance significantly above chance.

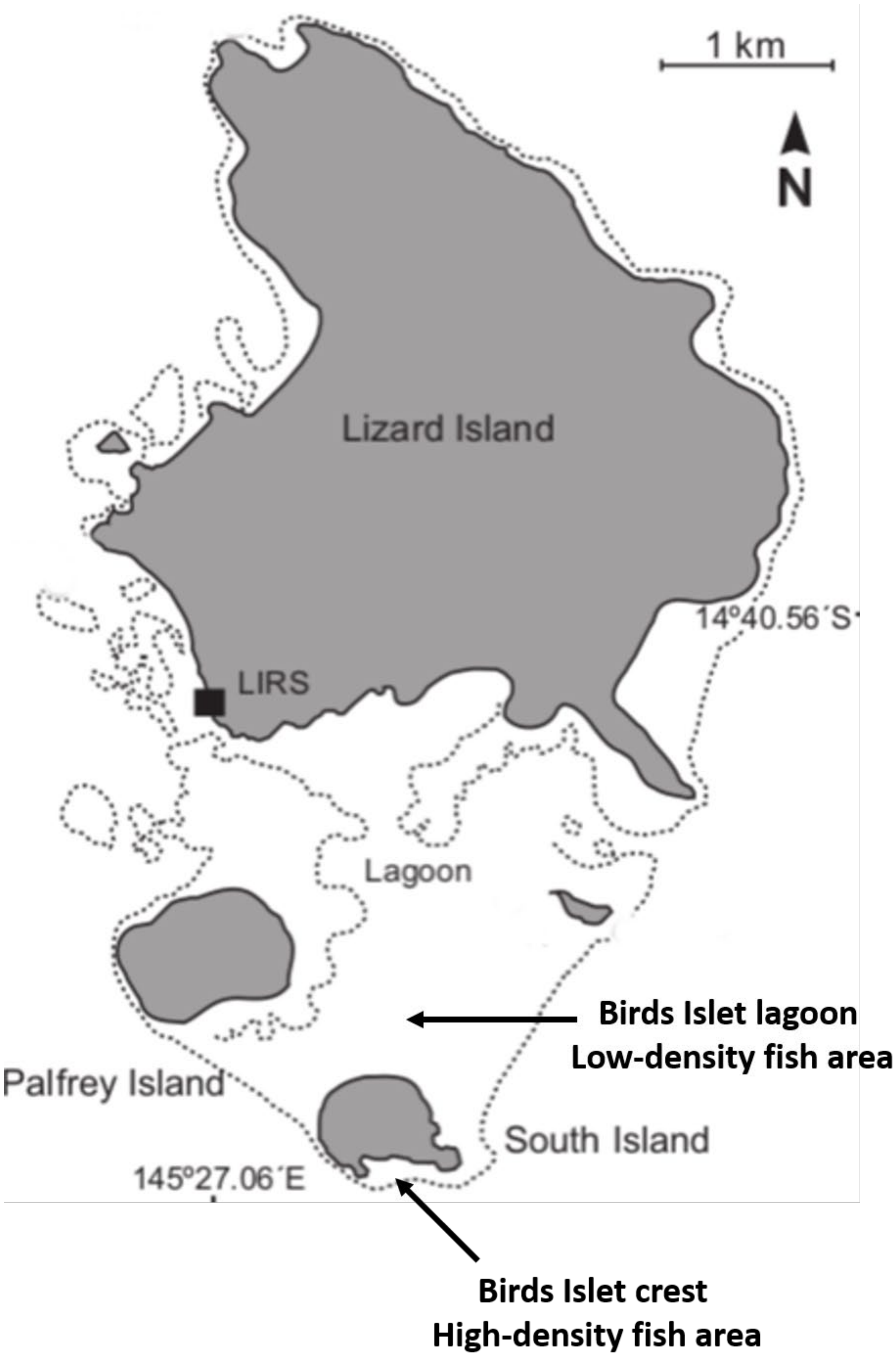


**S1 Fig: Study area showing the high-density and low-density fish sites.**

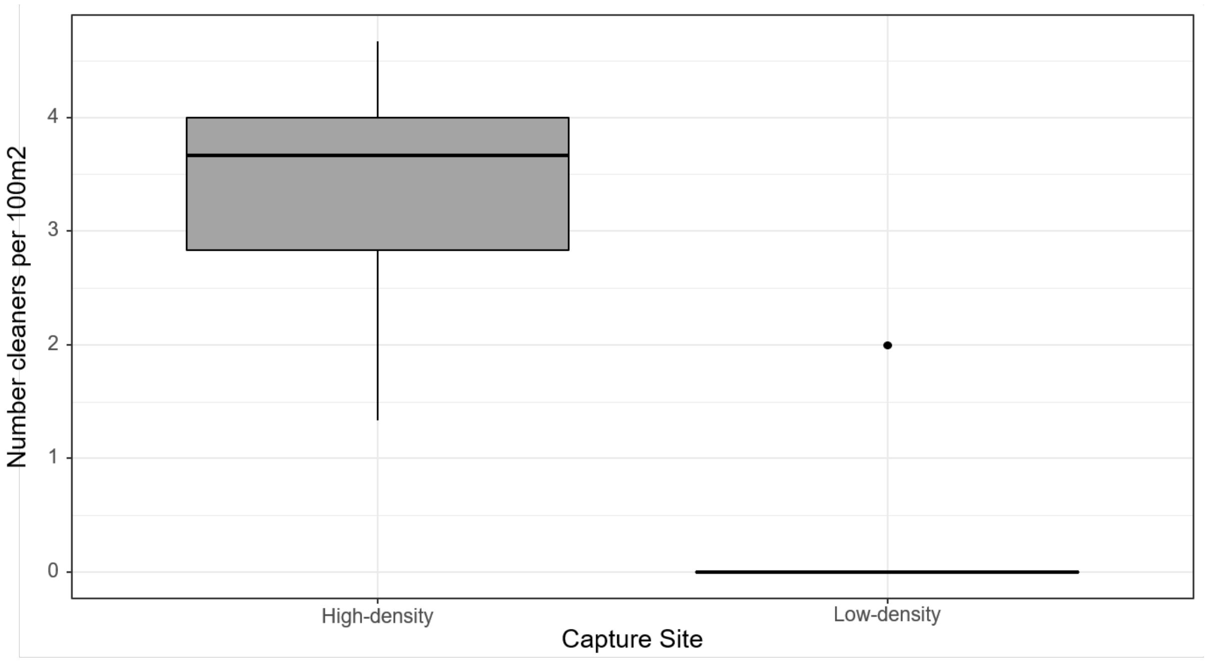


**S2 Fig: Box plot showing median, interquartiles and 95 % range of the number of cleaners per 100 m$^2$ split by site of capture (high-density and low-density site).**

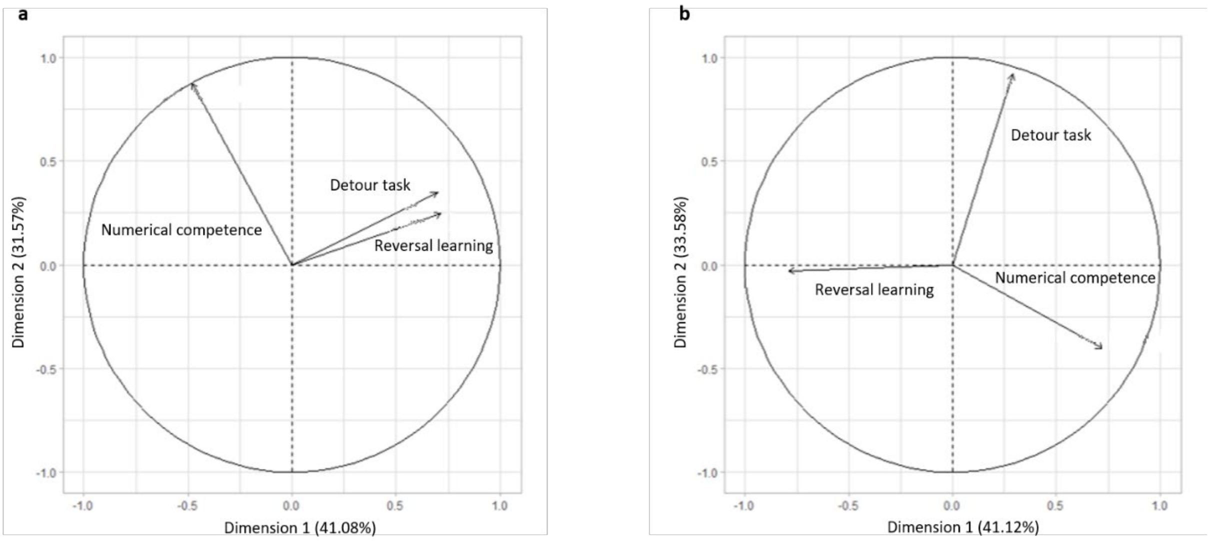


**S3 Fig: Principal Component Analysis (PCA) of the three cognitive tasks, separating data from individuals caught at the two different sites.** On the left (a): high fish density; dimension 1 (explaining 41.08 % of the variance in performance) and dimension 2 (explaining 31.57 % of the variance) are represented. On the right (b): low fish density; dimension 1 (explaining 41.12 % of the variance in performance) and dimension 2 (explaining 33.58 % of the variance). The results for each task are represented as vectors.

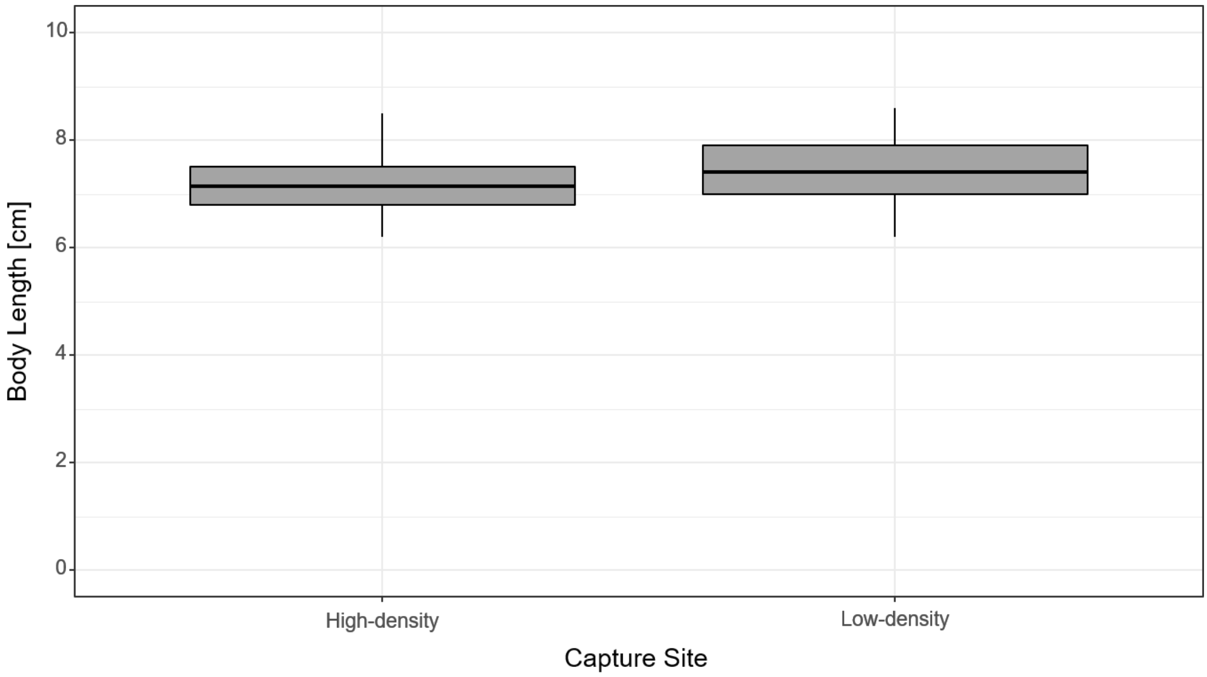


**S4 Fig: Box plot showing median, interquartiles and 95 % range of individual body length split by site of capture (high-density and low-density site).**

**The excluded task on object permanence: working memory (S5 Fig)**

Object permanence refers to the ability to understand that an object continues to exist even if it cannot be perceived directly at a given moment (1,2). To test for object permanence, an object of interest must be shown to a subject before it is moved out of sight. Subjects show evidence of object permanence if they are able to retrieve the object in a simultaneous choice task that offers also wrong options. Note that this task is not a learning task; individuals should be able to solve it spontaneously if they have object permanence.

We conducted three different versions of the task, invariably failing to obtain any evidence that cleaners can solve the task. The first version was implemented during the first field season with the first experimental set of fish. Three identical grey plates (4.5 cm x 4.5 cm) were inserted and placed 15 cm apart from each other in the experimental area. Two opaque barriers between the plates created three compartments (one for each plate; S5a,b Fig). Only one plate offered a visible food reward. After a few seconds, the plates were turned around. The fish had to choose the plate where the food reward was displayed. The setup was apparently intimidating many fish, as many did not approach the plates. Others immediately develop a side preference. As a consequence, we had either no data or performance at chance levels (33 % correct choices).

In a first attempt to make the task procedure less intimidating and potentially easier, we only used a single reward plate (one grey Plexiglas plate 4.5 cm x 4.5 cm with 2 green bands and 2 black dots). The plate was shown for a few seconds in either corner of the experimental compartment, before two identical barriers were placed in both corners to form two separate compartments (S5d Fig). Each barrier left a small open space for cleaners to enter the compartment. Cleaners had to choose the compartment hiding the plate in order to be allowed to eat the food; if they chose the wrong compartment, the reward plate was removed. We conducted 20 trials over 4 consecutive days, with the position of the plate counterbalanced within sessions of 10 trials. The cleaners' performance in this experiment was at chance levels (mean and SD of correct choices: 51.46 % ± 3.84).

In the second year, we attempted to make the information more salient for the cleaners. First, the two compartments in the two corners were in place before a trial started, as would be any reef structure in nature. Furthermore, we introduced the reward plate close to the holding compartment before moving it slowly to the end of the tank and let it slide out of sight into one compartment, like a client swimming out of sight on a reef (S5e,f Fig). Again, we conducted 20 trials over 4 consecutive days, with the position of the plate counterbalanced within sessions of 10 trials. The cleaners' performance in this experiment was again at chance levels (mean and SD of correct choices: 50.76 % ± 3.99). More generally, not a single individual ever deviated significantly from chance expectations in any of the three experimental setups. The task therefore did not measure object permanence in our sample, and was thus not included in the data. The data are publicly archived with the other data. To date, it is not clear whether these results show that cleaner fish lack object permanence. If so, this would suggest that their cognition is organized in an even more fundamentally different way than in vertebrates, not only lacking a positive manifold and thus *g.* However, since other fish apparently have object permanence (3), a valid alternative explanation of these results is that cleaner fish may well have object permanence, but that they did not succeed in this task for any non-cognitive reason (4).

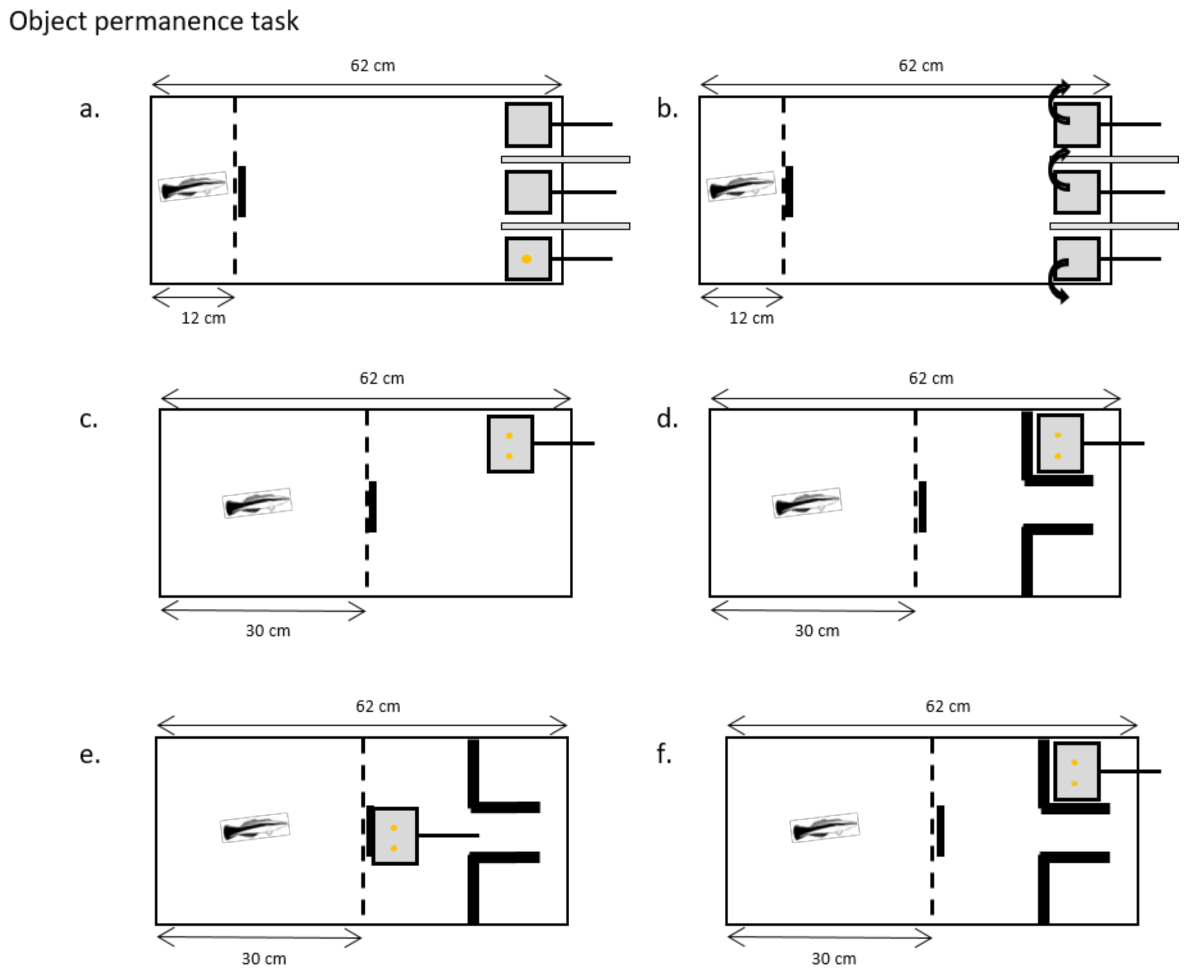


**S5 Fig. The three experimental sets to test for object permanence.** Dashed line: the transparent barrier that separated the holding compartment from the experimental compartment (with the short black line indicating the door through which fish could cross). Three plates were used the first set, while only one plate was used in the other two sets. Yellow dots signify a food item. The thick grey (a,b) and thick black (d-f) lines on the right sides of the aquaria represent opaque barriers. Panels a., c. and e. show how a trial was initially set up, while panels, b., d. and f. show the moment when the door was opened so that the fish could make its choice.
